# Activation of proneuronal transcription factor *Ascl1* in maternal liver ensures a healthy pregnancy

**DOI:** 10.1101/2021.04.27.441617

**Authors:** Joonyong Lee, Veronica Garcia, Shashank M. Nambiar, Huaizhou Jiang, Guoli Dai

**Affiliations:** Department of Biology, School of Science, Indiana University-Purdue University Indianapolis, Indianapolis, IN 46202

**Keywords:** *Ascl1*, hepatocyte, liver, pregnancy, insulin-like growth factor 2

## Abstract

**Background & Aims:** Maternal liver exhibits robust adaptations to pregnancy to accommodate the metabolic needs of developing and growing placenta and fetus by largely unknown mechanisms. We found that achaete-scute homolog-like 1 (*Ascl1*), a basic helix-loop-helix transcription factor essential for neuronal development, is highly activated in maternal hepatocytes during the second half of gestation in mice.

**Methods:** To investigate whether and how *Ascl1* plays a pregnancy-dependent role, we deleted the *Ascl1* gene specifically in maternal hepatocytes from mid-gestation until term.

**Results:** As a result, we identified multiple *Ascl1*-dependent phenotypes. Maternal livers lacking *Ascl1* exhibited aberrant hepatocyte structure, increased hepatocyte proliferation, enlarged hepatocyte size, reduced albumin production, and elevated release of liver enzymes, indicating maternal liver dysfunction. Simultaneously, maternal pancreas and spleen and the placenta displayed marked overgrowth; and the maternal ceca microbiome showed alterations in relative abundance of several bacterial subpopulations. Moreover, litters born from maternal hepatic *Ascl1*-deficient dams experienced abnormal postnatal growth after weaning, implying an adverse pregnancy outcome. Mechanistically, we found that maternal hepatocytes deficient for *Ascl1* exhibited robust activation of *insulin-like growth factor 2* expression, which may contribute to the *Ascl1*-dependent phenotypes widespread in maternal and uteroplacental compartments.

**Conclusion:** In summary, we demonstrate that maternal liver, via activating *Ascl1* expression, modulates the adaptations of maternal organs and the growth of the placenta to maintain a healthy pregnancy. Our studies reveal *Ascl1* as a novel and critical regulator of the physiology of pregnancy.

**Synopsis:** How the maternal liver adapts to pregnancy remains elusive. We found that maternal liver activates the expression of *Ascl1*, a gene encoding a proneuronal transcription factor, to coordinate the adaptations of maternal organs and the growth of the placenta, enabling a healthy pregnancy and normal postnatal growth of the offspring.

**Graphical Abstract:** 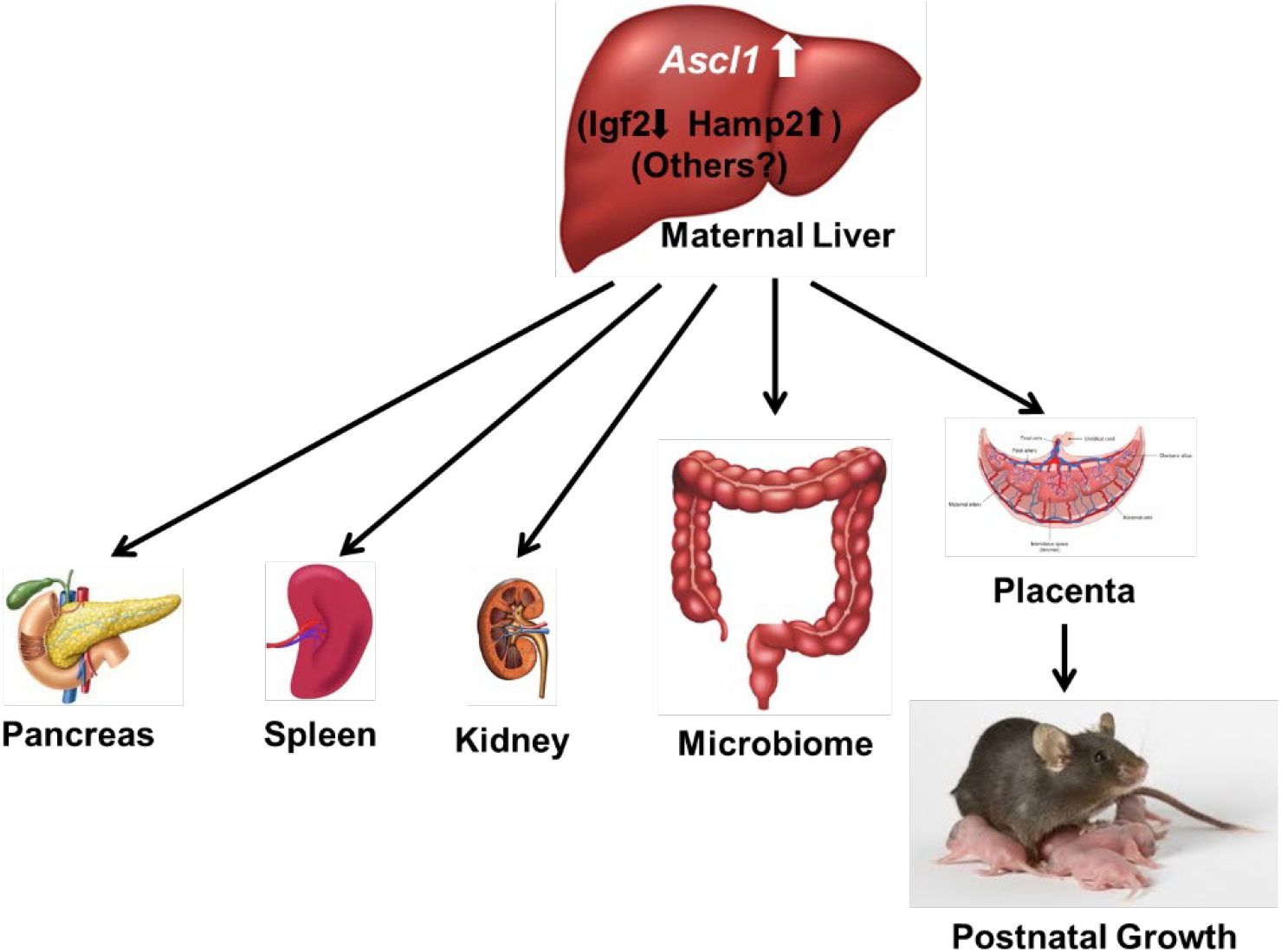

## Introduction

The establishment and maintenance of pregnancy requires highly coordinated adaptations in maternal, uteroplacental, and fetal compartments. As pregnancy progresses, in the maternal compartment, the liver grows to expand its metabolic capacity; the pancreas proliferates its β-cell population to elevate insulin production; the gut alters its microbiome contributing to immunological adjustments; the spleen undergoes development and growth of the erythroid lineage; and the kidney expands its volume allowing for increases in blood flow and fluid retention ^1–6^. These adaptations of maternal non-reproductive organs to pregnancy must be required to suffice the demands of the developing and growing placenta and fetus. However, the physiological significance and the regulatory mechanisms of these processes are poorly understood, representing an emerging research field.

Our work and that of others have demonstrated that maternal liver undergoes hyperplasia and hypertrophy, changes its gene expression profile, and thereby markedly expands during gestation in rodents ^7–9^. However, how this is controlled remains elusive. We previously showed that achaete-scute homolog-like 1 (*Ascl1*), a basic helix-loop-helix transcription factor essential for neurogenesis ^10–12^, is highly upregulated in maternal livers of pregnant rats ^13^. During development, *Ascl1* controls proliferation, cell cycle exit, and full neuronal differentiation and specification of neural progenitor cells in both the central and peripheral nervous systems ^11, 14, 15^ *Ascl1* inhibits its own expression by negative autoregulation in the developing nervous system, possibly explaining the lack of overt abnormalities in *Ascl1^+/-^* mice ^16^, whereas *Ascl1^+/−^* pups die within hours after birth due to defects in brain development ^10^. In the adult, ASCL1 is expressed only in the brain and spinal cord where there is ongoing neurogenesis, and in developing neuroendocrine cells in multiple organs including the cerebellum, thyroid, and thymus ^17^. In this study, we evaluated the activation, and explored the function, of *Ascl1* in maternal livers of pregnant mice. We found that *Ascl1* not only modulates maternal hepatic adaptation, but also mediates the communication of maternal liver with other maternal organs and the placenta, eventually affecting pregnancy outcomes.

## Materials and methods

### Mice

*Ascl1^fl/fl^*;*R26^EYFP/EYFP^* mice were a generous gift from Dr. Francois Guillemot (The Francis Crick Institute, Midland Road, London NW 1AT, UK) ^18^. The mice are referred hereafter as *Ascl1^fl/fl^* for simplicity. C57BL/6 mice were purchased from the Jackson Laboratory (Bar Harbor, ME, USA). Mice were maintained on a 12-hour light/12-hour dark cycle (7 AM on and 7 PM off) at 22 ± 1°C and given a standard rodent chow and water *ad libitum*. All of the animal experiments were conducted in accordance with the National Institutes of Health *Guide for the Care and Use of Laboratory Animals*. The protocols for the care and use of animals were approved by the Indiana University-Purdue University Indianapolis Animal Care and Use Committee.

### Mouse genotyping

Genomic DNA was prepared from mouse ear snips using the modified HotSHOT method ^19^. All mice were genotyped by polymerase chain reaction (PCR) using KAPA Taq PCR Kits (Kapa Biosystems, Inc., Wilmington, MA, USA). Specific primers purchased from Integrated DNA Technologies (Coralville, IA, USA) were used to detect the wild type and mutant alleles (Supplemental Table 1). Primers *Ascl1* Forward and *Ascl1* WT Reverse were used to detect the *Ascl1* wild type allele (342 bp), and primers *Ascl1* Forward and *Ascl1* Mutant Reverse were used to detect floxed *Ascl1* allele (857 bp) ^20^. PCR conditions were 35 cycles of 94°C/30 sec; 69°C/30 sec; 72°C/90 sec.

### Timed pregnancy and virus injection

Timed pregnancy was generated by mating 3-month-old virgin *Ascl1^fl/fl^* female mice with wild type male mice to ensure heterozygous fetuses with one wild type *Ascl1* allele. The presence of a copulation plug in the vagina was designated as gestation day (GD) 1. Adeno-associated virus serotype 8 (AAV8) with the thyroxine-binding globulin promoter (*TBG*) promoter expressing *Cre* (Addgene, AV-8-PV1091) were injected via tail vein at a dose of 1×10^12^ genomic copies per mouse on GD8, and mice were sacrificed on GD15 and GD18 using cervical dislocation. Adeno-associated viruses with a null vector (Addgene, AV-8-PV0148) were used as controls.

### Tissue collection and histology

Mice were euthanized at various time points. Maternal liver, pancreas, spleen, and kidney organs, and placentas and fetuses were collected and weighed. Part of each tissue was fixed in 10% neutral buffered formalin (NBF), embedded in paraffin and sectioned at 5 μm for hematoxylin and eosin (H&E) staining and histological analysis. Meanwhile, part of each tissue was embedded in optimal cutting temperature (OCT) compound (Fisher Scientific, 23-730-571) on heptane cooled in dry ice and stored at −80°C until processing. The remaining tissues were snap-frozen in liquid nitrogen and stored at −80°C for protein and RNA extraction.

### Immunohistochemistry

Formalin-fixed and paraffin-embedded (FFPE) maternal liver sections were subjected to standard immunohistochemistry. Primary antibodies against β-Catenin (BD Transduction, 610153, 1:100), IGF2 (R&D Systems, AF792, 1:1,000), and Ki67 (Thermo Fisher Scientific, RM-9106-S1, 1:100) were used for immunostaining. The slide images were acquired by the Leica DM2000 microscope using the Leica Application Suite program. β-Catenin-positive or Ki67-positive hepatocytes in five random fields of view at X400 magnification were counted using ImageJ software^21^. Periodic Acid Schiff staining was performed on FFPE placental sections in 0.5% periodic acid solution (Santa Cruz, sc-215695) and counterstained in hematoxylin (Leica, 3801575).

### Western blotting

Liver and placental homogenates (10-30 μg) were separated by polyacrylamide gel electrophoresis under reducing conditions and transferred to polyvinylidene difluoride (PVDF) membranes. The following antibodies were used: p-AKT (T308, abcam, ab76297, 1:5,000), p-AKT (S473, Epitomics, 2118-1, 1:2,000), ASCL1 (R&D Systems, BAF2567, 1:500), ERK1/2 (Cell Signaling, 9102, 1:2,000), p-ERK1/2 (T202/Y204, Cell Signaling, 4377, 1:2,000), GAPDH (Cell Signaling, 5174, 1:2,000), IGF2 (ABclonal, A2086, 1:1,000), Lamin B1 (Cell Signaling, 9087, 1:1,000), and PL-II (gift from Dr. Soares at the University of Kansas Medical Center, 1:2,000). Immune complexes were detected by SuperSignal West Pico PLUS Chemiluminescent Substrate (Thermo Fisher Scientific, 34577). Signals were detected using ImageQuant LAS 4000 Mini (General Electric Life Sciences) and quantified using ImageJ software ^21^.

### *In situ* hybridization

*In situ* hybridization was performed using RNAscope 2.5 HD Assay (Advanced Cell Diagnostics, 322310 and 322360) and appropriate probes and as per directions by the manufacturer. The following probes were used: *Ascl1* (Advanced Cell Diagnostics, 476321), *Igf2* (Advanced Cell Diagnostics, 437671), *PL-I* (Advanced Cell Diagnostics, 405521), and *PL-II* (Advanced Cell Diagnostics, 423681). A positive control probe *Ppib* (Advanced Cell Diagnostics, 310043) and a negative control probe *DapB* (Advanced Cell Diagnostics, 313911) were used to determine the efficacy of the protocol. These probes were applied to formalin-fixed and paraffin-embedded sections of mouse livers and placentas. The slide images were acquired by the Leica DM2000 microscope using the Leica Application Suite program. After approval from the institutional review boards, the laboratory information systems of our institute were searched. A set of paraffin blocks archived in the Department of Pathology of Indiana University School of Medicine were selected. They represented liver tissues from pregnant patients and patients with hepatocellular carcinoma or hepatocellular adenoma. These paraffin blocks were sectioned for *in situ* hybridization with a human Ascl1 probe (Advanced Cell Diagnostics, 459721).

### Quantitative real-time polymerase chain reaction (qRT-PCR)

Total RNA was isolated from snap-frozen liver tissue using TRIzol reagent (Invitrogen, 15596018) as per directions by the manufacturer. Complementary DNA (cDNA) was synthesized from 1 μg of total RNA using Verso cDNA kit (Thermo Fisher Scientific, AB1453B) and diluted four times with water. Quantitative real-time polymerase chain reaction (qRT-PCR) was performed using the diluted cDNA with either TaqMan Gene Expression Master Mix (Applied Biosystems, 4369016) or PowerUp SYBR Green Master Mix (Applied Biosystems, A25742) with specific gene probes **(Supplemental Tables 2 and 3)** ^22^. qRT-PCR was performed using the 7300 Real-Time PCR System (Applied Biosystems) and analyzed by the 7300 System SDS RQ Study Software (Applied Biosystems). qRT-PCR conditions for probes **(Supplemental Tables 2)** using TaqMan Gene Expression Master Mix were uracil N-glycosylase (UNG) incubation (50°C/2 min), polymerase activation (95°C/10 min), and 40 cycles of PCR (95°C/15 sec, 60°C/1 min). qRT-PCR conditions for probes **(Supplemental Tables 3)** using PowerUp SYBR Green Master Mix were uracil-DNA glycosylase (UDG) activation (50°C/2 min), polymerase activation (95°C/2 min), 40 cycles of PCR (95°C/15 sec, 60°C/15 sec, 72°C/1 min), and dissociation curve (95°C/15 sec, 60°C/30 sec, 95°C/15 sec). Relative gene expression was calculated by the comparative C_⊤_ method (ΔΔCt) and normalized to 18S rRNA transcript levels.

### RNA sequencing

Total RNA was isolated from snap-frozen liver tissue using the RNeasy Plus Mini Kit (Qiagen, 74134) as per directions by the manufacturer. RNA sequencing was performed and analyzed by the Center for Medical Genomics Core (Indiana University School of Medicine). In brief, total RNA concentration and quality were assessed using Agilent 2100 Bioanalyzer. Single-indexed strand-specific cDNA library from total RNA samples (500 ng input with RIN ≥ 5) was prepared using TruSeq Stranded mRNA Library Prep Kit (Illumina) as per directions by the manufacturer. The quality and size distribution of the libraries were assessed using Qubit and Agilent 2100 Bioanalyzer. Libraries (200 pM) were clustered and amplified on cBot using HiSeq 3000/4000 PE Cluster Kit and sequenced (2×75 bp paired-end reads) on HiSeq4000 (Illumina) using HiSeq 3000/4000 PE SBS Kit. A Phred quality score (Q score) was used to measure the quality of sequencing. The quality of the sequencing data was assessed using FastQC (Babraham Bioinformatics, Cambridge, UK). All sequenced libraries were mapped to the mouse genome (UCSC mm10) using STAR RNA-seq aligner, reads distribution across the genome was assessed using bamutils (from ngsutils), and uniquely mapped sequencing reads were assigned to mm10 refGene genes using featureCounts (from subread). Genes with CPM < 0.5 in more than 5 of the samples were removed. The data were normalized using the TMM method. Differential expression analysis was performed using edgeR and Ingenuity Pathway Analysis (IPA) with +I-2-fold change and P < 0.05. False discovery rate (FDR) was computed from p-values using the Benjamini-Hochberg procedure.

### Serum biochemistry

Blood from nonpregnant and pregnant mice (GD15 and GD18) was collected and left to clot at room temperature. After two centrifugations at 3000 RPM, serum was collected and analyzed by the Eli Lilly and Company (Indianapolis, IN). Serum insulin levels were measured by the Translation Core at the IU School of Medicine Center for Diabetes and Metabolic Diseases (Indiana University School of Medicine). Serum IGF2 levels were quantified using a 1/6 dilution with the Mouse IGF-2 ELISA Kit (Boster Biological Technology, EK0381) as per directions by the manufacturer and read with the SpectraMax M2e spectrophotometer using the SoftMax Pro 6 program.

### Microbiome 16S sequencing

Total microbial DNA was isolated from the snap-frozen cecal sample using the PureLink Microbiome DNA Purification Kit (Invitrogen, A29789) as per directions of the manufacturer. Microbiome 16S sequencing was performed and analyzed by the Zymo Research Corporation (Irvine, CA). In brief, bacterial 16S ribosomal RNA gene targeted sequencing was performed using the Quick-16S NGS Library Preparation Kit (Zymo Research, Irvine, CA). The bacterial 16S primers, custom-designed by Zymo Research, amplified the V3-V4 region of the 16S rRNA gene. The sequencing library was prepared using PCR reactions in real-time PCR machines to control cycles and therefore prevent PCR chimera formation. The final PCR products were quantified with qPCR fluorescence readings and pooled together based on equal molarity. The final pooled library was cleaned up with Select-a-Size DNA Clean & Concentrator (Zymo Research, Irvine, CA), then quantified with TapeStation and Qubit. The final library was sequenced on Illumina MiSeq with a v3 reagent kit (600 cycles). The sequencing was performed with >10% PhiX spike-in. Amplicon sequences were inferred from raw reads using the Dada2 pipeline. Chimeric sequences were also removed with the Dada2 pipeline. Taxonomy assignment, composition bar charts, alpha-diversity, and beta-diversity analyses were performed with Qiime v.1.9.1. Taxa that have an abundance significantly different among groups were identified by LEfSe with default settings if applicable. Differential expression analysis was assessed using IPA with +/- 2-fold change and *P* < 0.05.

### β-Cell mass

Pancreatic tissues were fixed in 4% paraformaldehyde (PFA) (Sigma Aldrich, 818715), embedded in paraffin, and sectioned at 7 μm for collecting 5 sections at 50 μm apart per pancreatic sample. Pancreatic sections immunostained for Insulin (Santa Cruz, sc-9168, 1:100) were used to quantify insulin-positive β-cell mass. β-Cell mass was determined by the ratio of the insulin-positive stained area to the tissue area multiplied by the pancreas weight.

### Statistical analysis

Data are shown as means ± standard deviation (SD) or means ± standard error of the mean (SEM). Statistical analyses were performed using a two-sided unpaired Student’s t-test with the means ± 95% confidence intervals. Significant differences were defined when *P* < 0.05.

## Results

### *Ascl1* is highly activated in maternal hepatocytes

We first evaluated the expression of *Ascl1* in maternal livers throughout the course of gestation in mice. As pregnancy advanced, its transcript levels in maternal livers were progressively elevated, reaching up to 26,000-fold on gestation days 13 and 15, relative to the pre-pregnancy state (Fig. 1A). Maternal hepatic *Ascl1* protein was abundantly expressed as its mRNA levels peaked on these two gestation days (Fig. 1B). *Ascl1* transcript was detected exclusively in maternal hepatocytes (Fig. 1C). In humans, *Ascl1* mRNA was abundantly expressed in a maternal liver of a pregnant woman, and in liver diseases such as hepatocellular carcinoma, hepatocellular adenoma, and hepatitis (Fig. 1D). Thus, it is truly astonishing that pregnancy activates a proneuronal transcription factor in an epithelial cell type in such a striking magnitude in a maternal compartment of pregnant animals and humans, strongly suggesting a gestation-dependent role for this transcription factor.

**Fig. 1.**
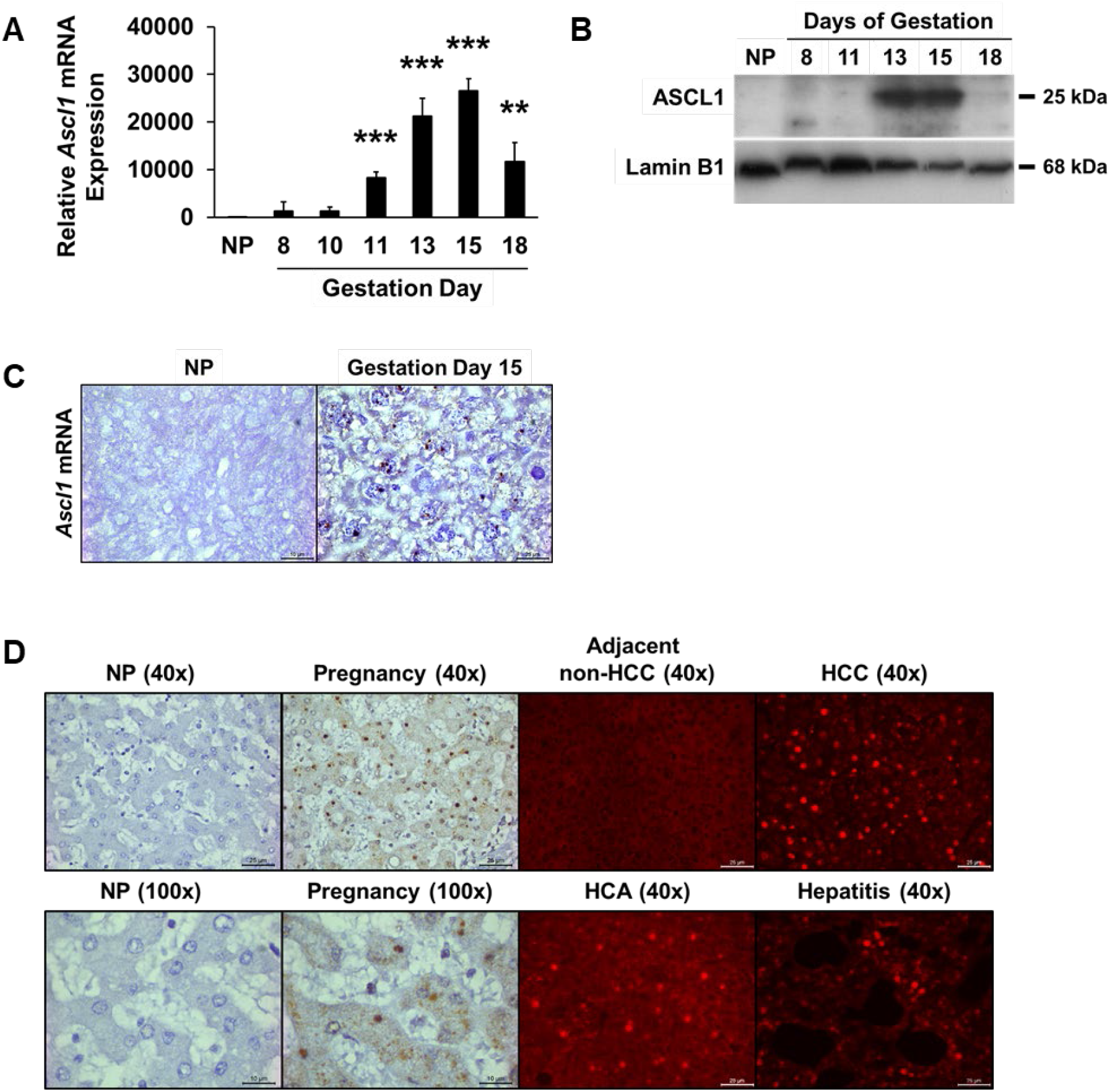
*Ascl1* activation and *Ascl1*-dependent transcriptome in maternal livers of pregnant mice. Timed pregnancies were generated in 3-month-old C57BL/6 female mice. Livers were collected from non-pregnant (NP) mice and maternal livers from pregnant mice at the indicated gestation days (GD). **(A-C)** *Ascl1* expression in maternal livers. **(A)** Hepatic *Ascl1* mRNA levels were measured using qRT-PCR and presented as the mean fold changes relative to NP controls ± SD (n = 3). **, *P* < 0.01; ***, *P* < 0.001, compared with NP controls. **(B)** Western blotting was performed using liver nuclear lysates with antibodies against ASCL1. Lamin B1 was used as a loading control. **(C)** Liver sections were subjected to *Ascl1 in situ* hybridization. *Ascl1* mRNA is stained dark brown. **(D)** Liver sections were prepared from archived paraffin blocks of human liver tissues. *Ascl1* mRNA were visualized on liver sections. Representative results are shown with liver sections from a nonpregnant (NP) woman (a 6-month pregnant mother), a hepatocellular carcinoma (HCC) patient, a hepatocellular adenoma (HCA) patient, and a 4.5-month child with hepatitis.

### Adeno-associated virus 8 (AAV8) containing the gene for *Cre* recombinase under the control of hepatocyte-specific thyroxine-binding globulin (*TBG*) promoter (AAV8-TBG-Cre) efficiently deletes *Ascl1* in maternal hepatocytes

*Ascl1* is activated in maternal liver during the second half of pregnancy, suggesting a role in the maintenance of pregnancy. To determine this, we took a loss-of-function approach to delete the *Ascl1* gene specifically in maternal hepatocytes from mid-gestation to term and evaluated how pregnancy was affected. We generated timed pregnancies in *Ascl1^fl/fl^* female mice by mating them with *Ascl1^+/+^* males. This led to a homogeneous *Ascl1^+/fl^* fetal genotype. AAV8-TBG-Cre virus or AAV8-TBG-null (control) virus was injected into *Ascl1^fl/fl^* mice on gestation day 8. Mice were sacrificed on gestation days 15 and 18 for phenotypic assessment. This AAV8-TBG virus has been shown to specifically infect hepatocytes with >99% efficiency in mice ^23, 24^. qRT-PCR analysis showed that *Ascl1* mRNA was almost completely lost in maternal livers of pregnant animals treated with the AAV8-TBG-Cre virus (Fig. 2A), indicating highly efficient *Ascl1* gene deletion in this organ. As such, mice deficient for *Ascl1* in maternal hepatocytes were referred to as hep-*Ascl1*^-/-^ mice hereafter. Of note, in a pretest study, we injected AAV8-TBG-GFP reporter virus into pregnant mice. As a result, we detected GFP expression in maternal livers but not in fetal livers (data not shown). This indicates that the AAV8 virus does not effectively pass through the placenta to infect the fetuses.

**Fig. 2.**
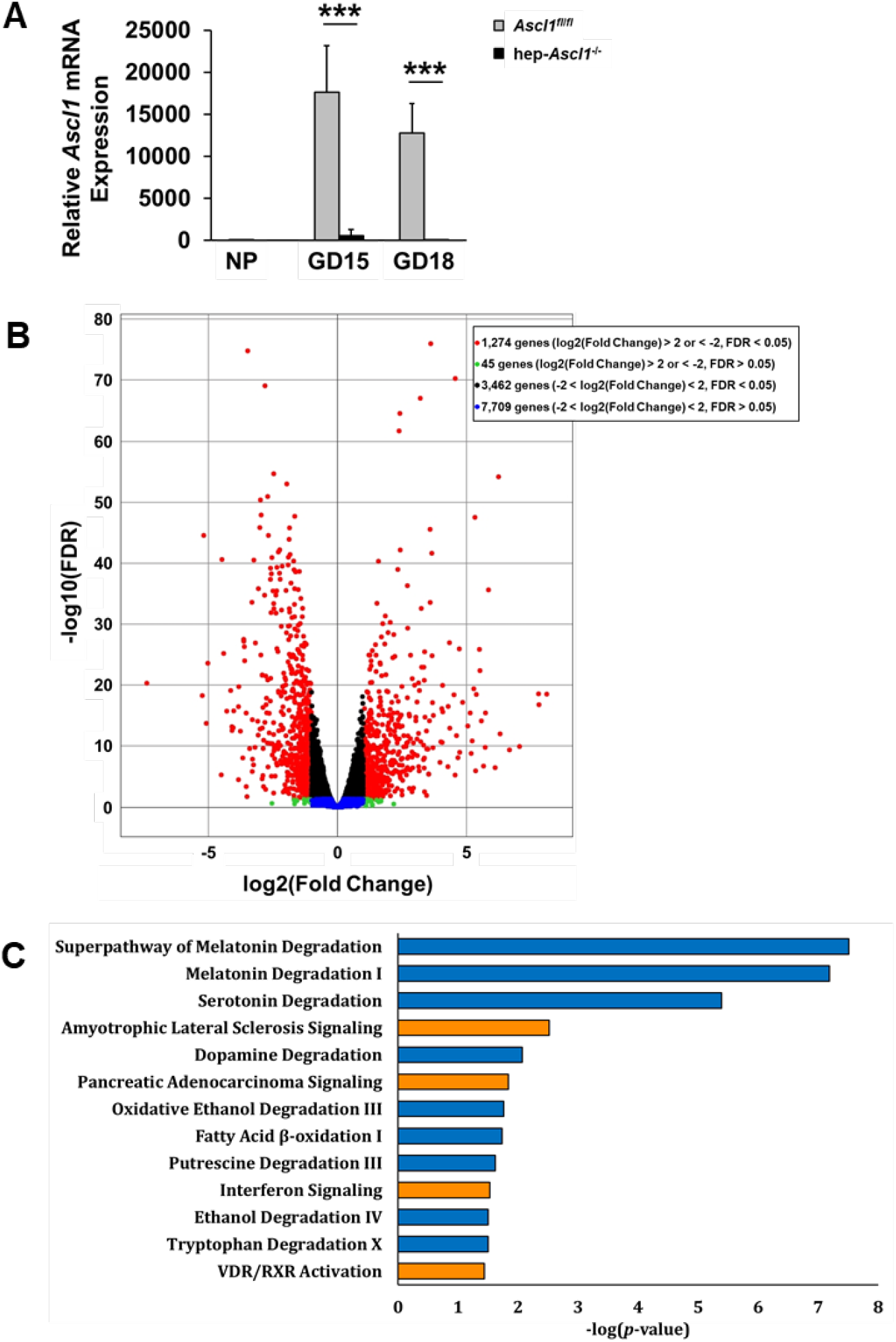
Differentially expressed genes and canonical pathways affected by hepatocyte-specific deletion of *Ascl1* in the maternal liver. **(A)** Total RNA was isolated from livers of nonpregnant (NP) and pregnant *Ascl1^fl/fl^* and hepatocyte-specific Ascl1 knockout (*hep-Ascl1^-/-^*) mice. Hepatic *Ascl1* mRNA levels were measured using qRT-PCR and presented as the mean fold changes relative to NP controls ± SD (n = 4-5). ***, *P* < 0.001. **(B-C)** *Ascl1*-dependent transcriptome. Total RNA was isolated from livers of *Ascl1^fl/fl^* and *hep-Ascl1^-/-^* mice on gestation day (GD) 15 and was subjected to RNA-sequencing (n = 4-5). **(B)** Differentially expressed genes are presented by the volcano plot. Red: significantly up- or down-regulated genes; black, green, and blue: non-significant genes. Differentially expressed genes with at least two-fold and *P* < 0.05 were analyzed using the Ingenuity Pathway Analysis (IPA). **(C)** Top enriched canonical pathways targeted by hepatic *Ascl1* are presented. Orange, up-regulated; blue, down-regulated.

### RNA-sequencing (RNA-seq) analysis reveals *Ascl1*-dependent transcriptome in maternal

We next compared the transcriptomes of gestation day 15 maternal livers between the two genotype groups of mice by RNA-seq to profile potential Ascl1 target genes in this context. This revealed 1,274 differentially expressed genes. They were either up-regulated or down-regulated by at least 2-fold with a false discovery rate of less than 0.05 when maternal hepatic *Ascl1* was lacking (Fig. 2B). Pathway analysis suggests that these genes are associated with many biological processes including the metabolism of hormones (e.g. melatonin), neurotransmitters (e.g. serotonin and dopamine), fatty acids, alcohol, carbohydrates, nucleic acids, and amino acids, cell cycle control, cell function and maintenance, and vitamin D receptor/retinoid X receptor activation (Fig. 2C). These data imply that *Ascl1* directly or indirectly regulates a broad spectrum of genes and thereby possesses multiple functions in maternal liver in this physiological state (pregnancy).

### Hepatocyte-specific *Ascl1* knockout results in maternal liver abnormalities

We subsequently assessed whether maternal livers exhibited *Ascl1*-dependent phenotypes. Compared with maternal hepatocytes sufficient for *Ascl1*, maternal hepatocytes lacking *Ascl1* displayed an aberrant structure (eosin staining-negative around the nuclei) (Fig. 3A), enhanced hepatocyte proliferation (Fig. 3B, Supplemental Fig. 1A), and increased size (Fig. 3C, Supplemental Fig. 1B), hence causing further enlargement of the maternal livers (Fig. 3D-E). These abnormalities were accompanied by reduced albumin mRNA expression and protein production and elevated circulating alanine transaminase (ALT) and aspartate aminotransferase (AST) (Fig. 3F). These observations indicate that *Ascl1* loss of function in maternal hepatocytes impairs their structure and function.

**Fig. 3.**
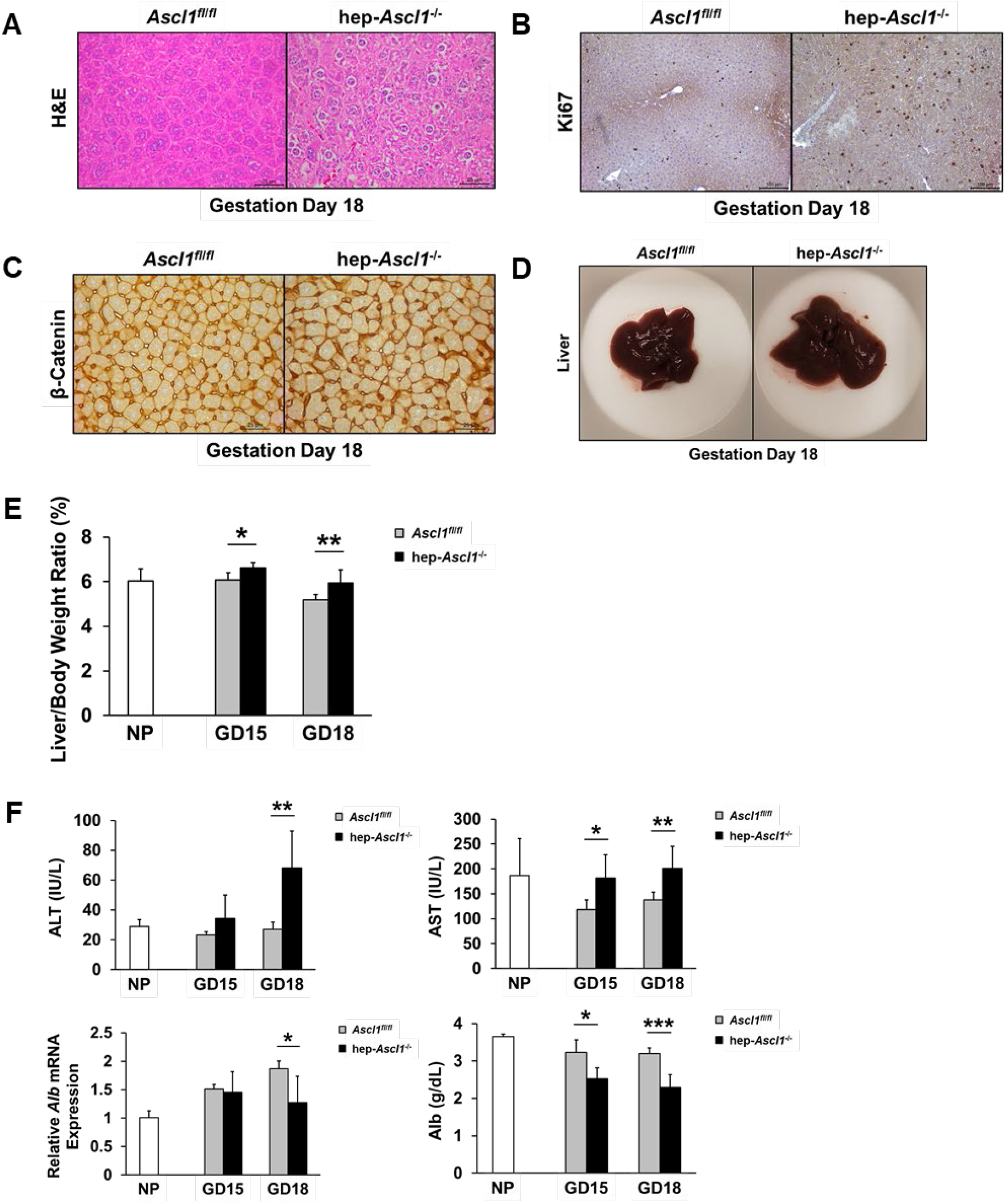
Maternal liver phenotypes in maternal hepatocyte-specific *Ascl1* ablated mice. Livers were collected and weighed from nonpregnant (NP) and gestation days (GD) 15 and 18 *Ascl1^fl/fl^* and *hep-Ascl1*^-/-^ mice. **(A)** Liver sections were stained with hematoxylin and eosin (H&E). Liver sections were subjected to **(B)** Ki67 and **(C)** β-Catenin staining. **(D)** Morphology of *Ascl1^fl/fl^* and *hep-Ascl1^-/-^* mouse livers. **(E)** Liver-to-body weight ratios are presented as means ± SD (n = 4-10). **(F)** Serums were collected from NP and pregnant *Ascl1^fl/fl^* and hep-*Ascl1*^-/-^ mice. Data of serum biochemical profile are expressed as means ± SD (n = 5). Hepatic *Albumin* mRNA levels were measured using qRT-PCR and presented as the mean fold changes relative to NP controls ± SD (n = 4-5). *, *P* < 0.05; **, *P* < 0.01; ***, *P* < 0.001.

### Hepatocyte-specific *Ascl1* knockout causes overgrowth of maternal pancreas, spleen, and kidney

We additionally examined several other maternal organs to estimate whether there were *Ascl1-*dependent systemic effects in the maternal compartment. Surprisingly, without maternal hepatic *Ascl1*, the maternal pancreas nearly doubled in size (Fig. 4A-B), had a reduced proportion of insulin-positive areas (Fig. 4C-D), and unchanged total β-cell mass (Fig. 4E), while the concentrations of circulating insulin and non-fasting blood glucose were unaltered (Supplemental Fig. 2). These data suggest that *Ascl1* inactivation in maternal liver stimulates the expansion of the exocrine component without interfering with the pregnancy-dependent growth of the endocrine component in the maternal pancreas. Similarly, the maternal spleen also almost doubled in volume with overtly expanded red and while pulps (Fig. 5A-C), while the maternal kidney showed enlargement without an obvious histological alteration (Fig. 5D-E). Collectively, we conclude that the loss of function of *Ascl1* in maternal hepatocytes imposes systematic effects, causing overgrowth of at least a subset of maternal organs.

**Fig. 4.**
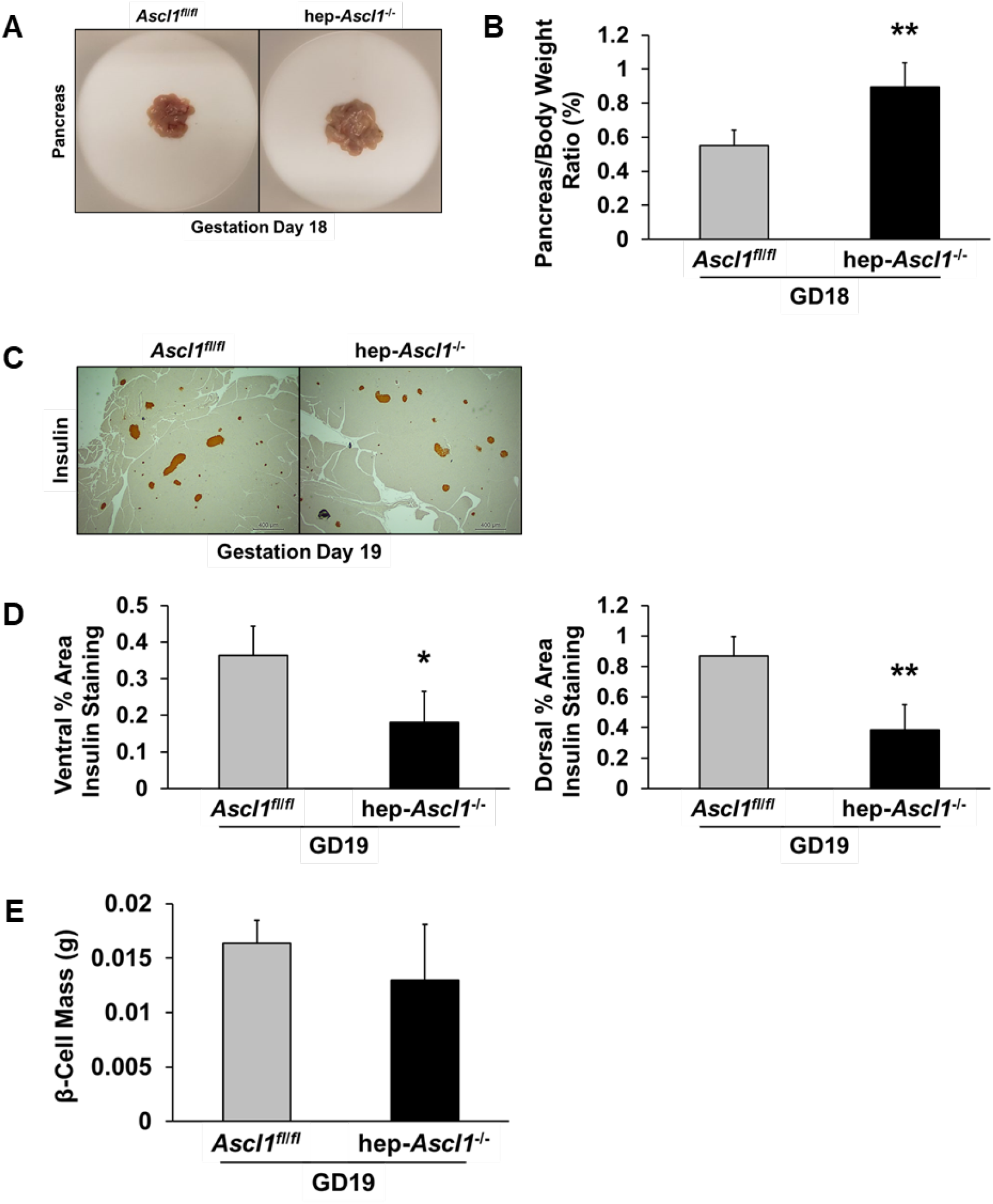
Maternal pancreas phenotypes in maternal hepatocyte-specific *Ascl1* ablated mice. Maternal pancreases were collected and weighed from nonpregnant (NP) and gestation day (GD) 18 *Ascl1^fl/fl^* and *hep-Ascl1^-/-^* mice. **(A)** Morphology of the pancreas. **(B)** Pancreas-to-total body weight ratios are presented as means ± SD (n = 4-5). **(C)** Pancreatic sections were subjected to insulin immunostaining. **(D)** The quantification of the insulin-positive islets to the pancreatic section area are presented as means ± SD (n = 4-5). **(E)** β-Cell mass is presented as the means ± SD (n = 4-5). *, *P* < 0.05; **, *P* < 0.01.

**Fig. 5.**
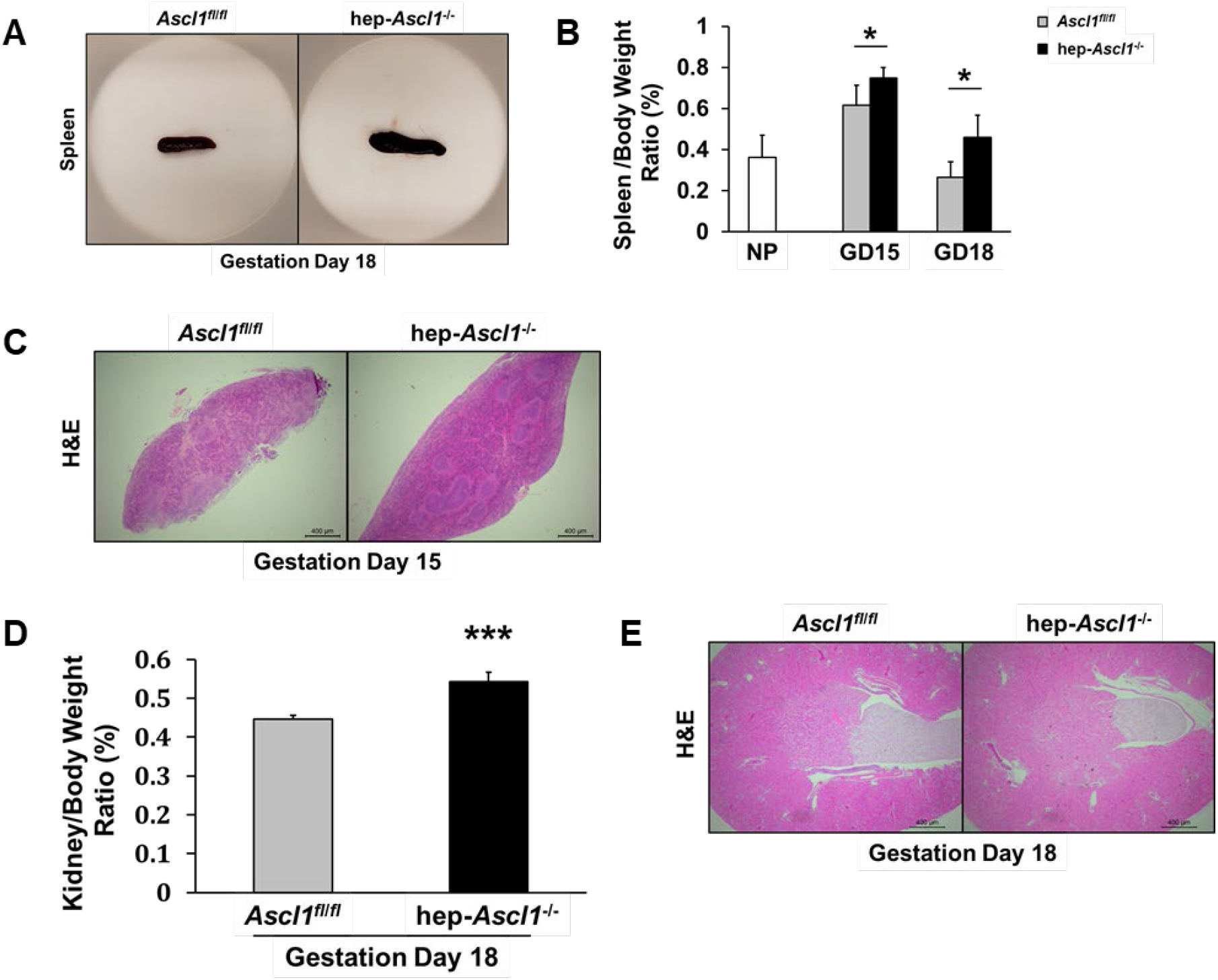
Maternal spleen and kidney phenotypes in maternal hepatocyte-specific *Ascl1* ablated mice. Maternal spleens and kidneys were collected and weighed from nonpregnant (NP) and gestation days (GD) 15 and 18 *Ascl1^fl/fl^* and *hep-Ascl1^-/-^* mice. **(A)** Morphology of the spleens. **(B)** Spleen-to-total body weight ratios are presented as means ± SD (n = 4-10). **(C)** Splenic sections were stained with hematoxylin and eosin (H&E). **(D)** Kidney-to-total body weight ratios are presented as means ± SD (n = 4-5). **(E)** Kidney sections were stained with H&E. *, *P* < 0.05; ***, *P* < 0.001.

### Hepatocyte-specific *Ascl1* knockout leads to abnormal maternal cecal microbiota

It is known that the maternal gut microbiome undergoes pregnancy-dependent changes, which are associated with the health of both the mother and the fetus ^25^. To determine whether maternal hepatic *Ascl1* is relevant to the maternal microbiome, we collected maternal cecal contents from gestation day 18 hep-*Ascl1*^-/-^ mice and *Ascl1^fl/fl^* (control) mice, and compared their microbiome profiles via 16S sequencing (Fig. 6A). Most notably, *Ascl1* loss in maternal hepatocytes resulted 1) in a complete depletion of *Pseudobutyrivibrio-Roseburia intestinalis*, which protects colonic mucosa against inflammation ^26^, 2) in the appearance of *Desulfovibrio oxamicus-vulgaris*, which metabolizes a variety of chemicals ^27^, and 3) in alterations in the relative abundance of 6 other bacteria species. Furthermore, RNA-seq analysis revealed that *Hamp2*, a gene encoding hepcidin antimicrobial peptide 2, which has a strong antimicrobial activity against certain bacteria ^28, 29^, depends on Ascl1 for expression. This finding was confirmed by qRT-PCR (Fig. 6B). Together, we demonstrate that maternal hepatic *Ascl1* modulates the adaptation of the maternal microbiota to pregnancy, potentially via regulating *Hamp2*.

**Fig. 6.**
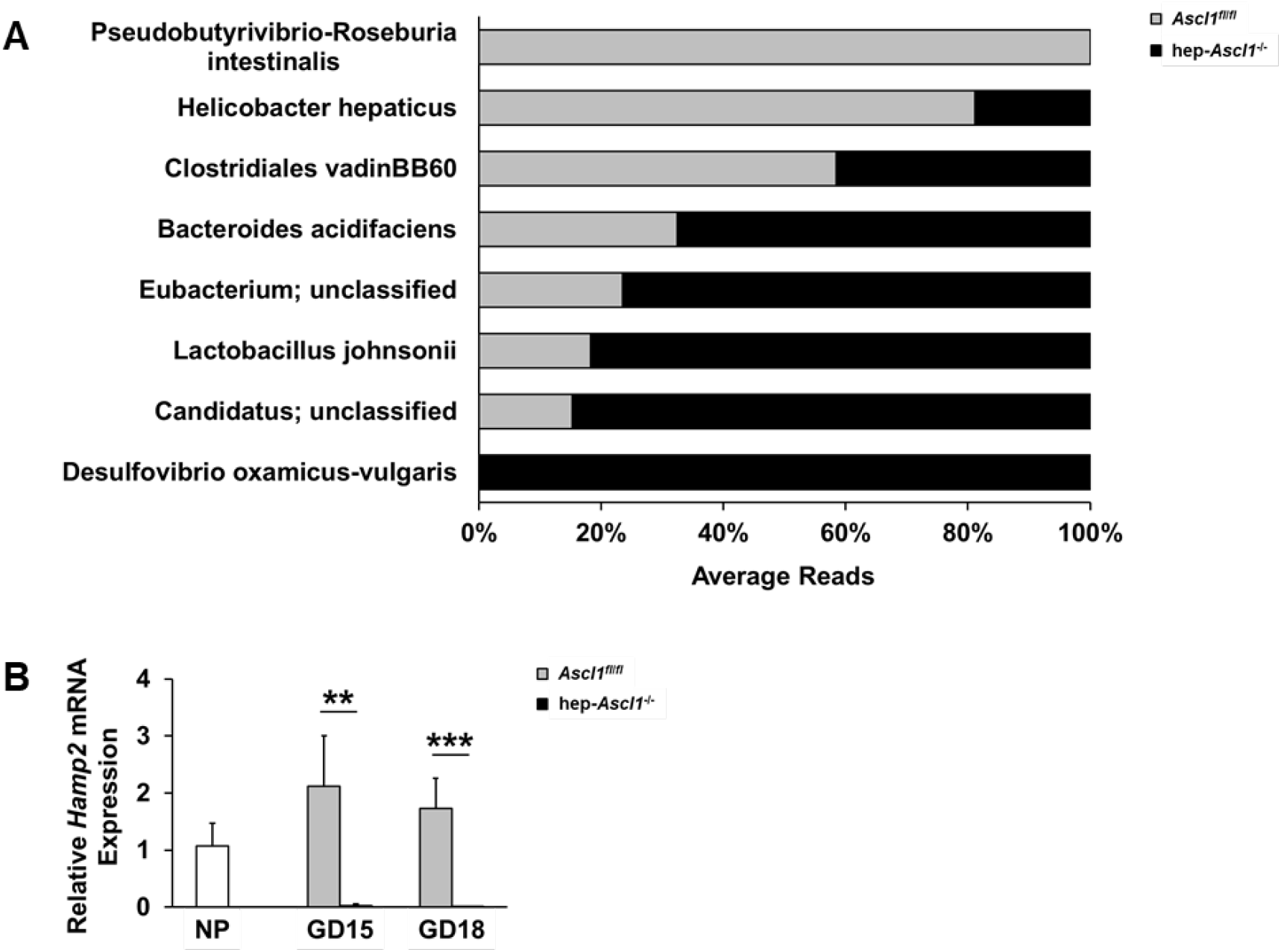
Phenotypes of maternal cecal microbiota in maternal hepatocyte-specific *Ascl1* ablated mice. Maternal cecal samples were collected from gestation day (GD) 18 *Ascl1^fl/fl^* and *hep-Ascl1^-/-^* mice. **(A)** Cecal microbiota analysis (n = 5). **(B)** Hepatic hepcidin antimicrobial peptide 2 (*Hamp2*) mRNA levels were measured using qRT-PCR and presented as the mean fold changes relative to nonpregnant (NP) controls ± SD (n = 4-5). **, *P* < 0.01; ***, *P* < 0.001.

### Hepatocyte-specific *Ascl1* knockout causes placental overgrowth and aberrant postnatal growth of offspring

We examined the uteroplacental and fetal compartments and postnatal growth of pups to determine whether *Ascl1* deficiency in maternal hepatocytes ultimately affects pregnancy outcomes. Compared to the placentas of control mice, the placentas of hep-*Ascl1^-/-^* mice were markedly enlarged, manifested by a 26.9% increase in weight on gestation day 15 and 33% on gestation day 18, with the expansion of both the junctional and labyrinth zone (Fig. 7A-B). The distribution of glycogen trophoblast cells visualized by glycogen staining did not appear to be *Ascl1*-dependent (Supplemental Fig. 3A). By *in situ* hybridization, we probed the mRNAs of *placental lactogen (PL)-I*, a marker gene for parietal trophoblast giant cells ^30^, and *PL-II*, a marker gene for parietal trophoblast giant cells, spongiotrophoblast cells, and labyrinth trophoblast giant cells ^30^, and did not observe an overt *Ascl1*-dependent distribution of these trophoblast cell populations (Supplemental Fig. 3B). These results of placental structural evaluations suggest that the loss of function of *Ascl1* in maternal hepatocytes causes placental overgrowth without disrupting placental structure. Moreover, when comparing hep-*Ascl1*^-/-^ pregnant mice with their control mice, the placental levels of insulin-like growth factor (IGF2), a potent placental and fetal growth factor ^31–33^, were reduced by 32% on gestation day 15 but were equivalent on gestation day 18, whereas placental concentration of PL-II, a major hormone produced by the placenta during the second half of gestation ^34^, was unchanged (Fig. 7C, Supplemental Fig. 3C). However, when maternal hepatic *Ascl1* was deficient, maternal blood concentration of alkaline phosphatase, an enzyme primarily elaborated from the placenta and a marker for trophoblast differentiation ^35^, was significantly elevated on gestation day 15 and more than 2-fold on gestation day 18. The data of these functional assessments suggest that *Ascl1* inactivation in maternal hepatocytes partially influences placental function.

**Fig. 7.**
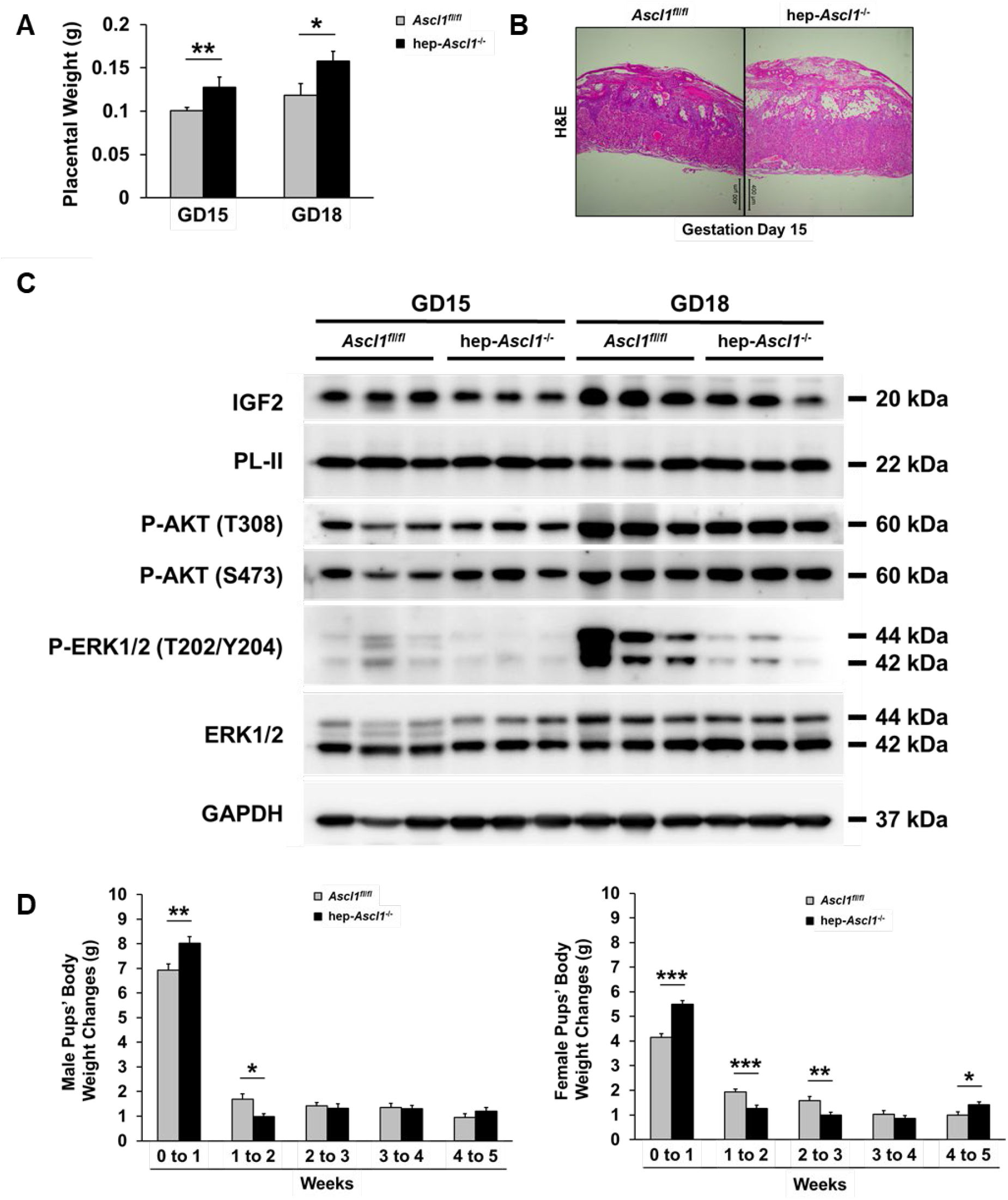
Placental phenotypes in maternal hepatocyte-specific *Ascl1* ablated mice. Placentas were collected and weighed from gestation days (GD) 15 and 18 *Ascl1^fl/fl^* and *hep-Ascl1^-/-^* mice. **(A)** Placental weights are presented as means ± SD (n = 4-9 dams). **(B)** Placental sections were stained with hematoxylin and eosin (H&E). **(C)** Western blotting was performed using placental lysates prepared from one placenta per dam with antibodies against the proteins indicated. Glyceraldehyde 3-phosphate dehydrogenase (GAPDH) was used as a loading control. **(D)** Weekly body weight changes of weaned pups (n = 43 from 5 *Ascl1^fl/fl^* dams; n = 44 from 6 *hep-Ascl1^-/-^* dams; n = 6-10 per litter) are presented as means ± SEM. *, *P* < 0.05; **, *P* < 0.01; ***, *P* < 0.001.

Furthermore, it is known that AKT and ERK signaling critically regulates placental development and growth ^36, 37^. We found that, without *Ascl1* in maternal hepatocytes, placental AKT1 phosphorylation at S473, but not at T308, was mildly increased; in contrast, ERK1 and ERK2 activities were dramatically inhibited, reduced as much as 80% and 76% respectively, prior to parturition (gestation day 18). Thus, the loss of maternal hepatic *Ascl1* results in strong inhibition of placental ERK signaling.

Additionally, we did not observe a difference in fetal weight and number on both gestation days 15 and 18 between the two genotype groups of mice (Supplemental Fig. 4). However, compared to those male and female pups born from control dams, male pups born from hep-*Ascl*^-/-^ dams displayed a 16% increase in weekly body weight gain at week 1 post-weaning, but a 41% decrease at week 2, while females pups born from hep-*Ascl1*^-/-^ dams exhibited a 32% increase at week 1, a 35% decrease at week 2, a 38% decrease at week 3, and a 43% increase at week 5 (Fig. 7D). Apparently, pups born from the two genotype groups of dams grew quite differently after weaning. Taken together, we demonstrate that the maternal hepatic *Ascl1* null mutation causes placental overgrowth, partial changes in placental function, severe suppression of placental ERK activity, and an abnormal postnatal growth pattern of offspring.

### Hepatocyte-specific *Ascl1* knockout leads to *Igf2* activation in maternal hepatocytes

The above observations that several maternal organs were significantly enlarged in hep-*Ascl1*^-/-^ pregnant mice relative to controls prompted us to look for growth factors regulated by *Ascl1* from our RNA-seq analysis. We found one candidate: *Igf2*, which encodes a potent growth factor ^31–33^ and was induced with Ascl1 loss. To validate the RNA-seq data, we quantified *Igf2* mRNA levels by qRT-PCR in maternal livers of gestation days 15 and 18 and show that the loss of *Ascl1* in maternal hepatocytes induced robust *Igf2* mRNA expression in maternal livers, most strikingly on gestation day 18 (Fig. 8A). Consistently, maternal hepatic IGF2 protein was abundantly produced in hep-*Ascl*^-/-^ pregnant mice, but was undetectable in control pregnant mice (Fig. 8B). *In situ* hybridization and immunohistochemistry detected rich *Igf2* transcripts (Fig. 8C) and IGF2 protein (Fig. 8D) in maternal hepatocytes deficient for *Ascl1*, but not in controls (Fig. 8D). In mice, *Igf2* is maternally imprinted and is differentially regulated in the placenta and fetus. It is transcribed from four promoters, designated *Igf2-P0, P1, P2, and P3. Igf2-P0* directs *Igf2* transcription in the placenta, whereas *Igf2-P1, P2, and P3* direct its transcription in both the placenta and fetus ^32, 38^. We used a set of qRT-PCR primers to analyze promoter-specific *Igf2* transcripts in maternal livers ^39^ (Fig. 8E).

We found that *P0* remained silent no matter the presence or absence of *Ascl1* in maternal hepatocytes. In contrast, the other three promoters were all activated due to *Ascl1* loss in these cells. The concentration of circulating IGF2 protein in gestation day 18 *hep-Ascl1^-/-^* mice was about 9 times higher than that in control mice (Fig. 8F). These data collectively demonstrate that, in maternal hepatocytes, *Ascl1* deficiency results in *Igf2* activation via *P1*, *P2*, and *P3*, creating a maternal environment rich in this growth factor. Thus, here we linked *Igf2* to *Ascl1*-dependent phenotypes in our experimental settings.

**Fig. 8.**
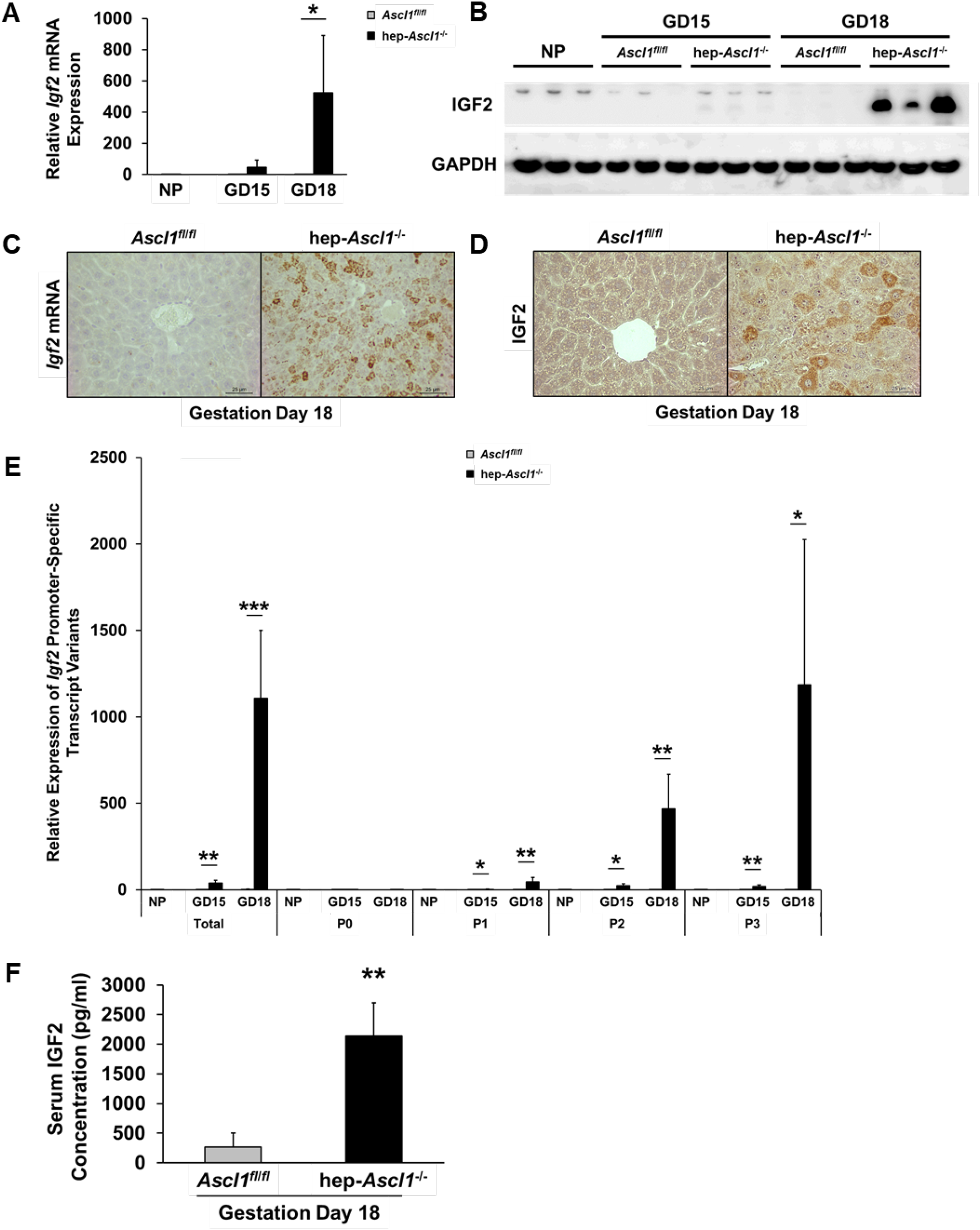
*Igf2* activation in maternal livers deficient for *Ascl1*. Maternal livers were collected and weighed from nonpregnant (NP) and gestation days (GD) 15 and 18 *Ascl1^fl/fl^* and *hep-Ascl1^-/-^* mice. **(A)** Hepatic *Igf2* mRNA levels were measured using qRT-PCR and presented as the mean fold changes relative to NP controls ± SD (n = 4-5). **(B)** Western blotting was performed using liver lysates with an antibody against IGF2. **(C)** *Igf2 in situ* hybridization on liver sections. **(D)** IGF2 immunostaining. **(E)** Levels of hepatic *Igf2* promoter-specific transcript variants were measured using qRT-PCR and presented as the mean fold changes relative to NP controls ± SD (n = 4-5). **(F)** Levels of IGF2 protein in serum were measured using ELISA and presented as the mean fold changes ± SD (n = 3-5). *, *P* < 0.05; **, *P* < 0.01; ***, *P* < 0.001. P0, placental-specific *Igf2* promoter; P1-3, placental- and fetal liver-specific *Igf2* promoter.

## Discussion

It has been an enigma why maternal liver activates a proneuronal gene (*Ascl1*) in response to pregnancy. Here we demonstrate that *Ascl1* is required for the maternal liver to structurally and functionally adapt to this physiological stimulus (pregnancy). Loss of *Ascl1* in maternal hepatocytes impairs their cellular structure, reflected by the formation of a layer of eosin staining-negative substance around their nuclei. This substance was Oil Red O staining- and Periodic acid Schiff staining-negative, suggesting that it is not a fatty acid or glycogen (data not shown). Others reported a similar phenomenon in *Mst1/2* double knockout hepatocytes ^40^. The connection between Ascl1 and Mst1/2 signaling in maternal hepatocytes will be interesting to explore in future. Although the eosin staining-negative substance around the hepatocyte nuclei remains unknown, we believe that it is linked to *Ascl1* deficiency-caused dysfunction of maternal hepatocytes, characterized by reduced albumin production and increased ALT and AST release. In addition, *Ascl1* loss induced increased proliferation and hypertrophy of maternal hepatocytes, which was associated with increased IGF2 production by these cells. IGF2 is a prototypical growth factor driving cell expansion and organogenesis during development ^41^, and adult regenerating livers activate IGF2 to promote hepatocyte replication ^42^. Here we show that pregnancy reprograms maternal hepatocytes to activate *Ascl1*, thereby controlling the transcriptional output and modulating the expression of at least 1,274 genes. Of special note, the top canonical pathway describing these differentially expressed genes is one that represents genes involved in the metabolism of neurotransmitters. *Ascl1* is known to be essential for the development of serotonergic and dopaminergic neurons and also regulates their neurotransmitter biosynthesis ^43, 44^. Remarkably, *Ascl1* alone is sufficient to generate functional neurons from fibroblasts and embryonic stem cells, being a key driver of induced neuronal cell reprogramming in different cell contexts ^45^. These findings raise the question of whether *Ascl1* activation enables maternal hepatocytes to possess some properties of neurons as an adaptive response to support a healthy pregnancy. We are very interested in answering this question in our future studies.

It is surprising that inactivating *Ascl1* in maternal liver induced robust systemic responses in the maternal compartment. This implies that the maternal liver, via activating *Ascl1*, communicates with other maternal organs to systematically coordinate maternal adaptations to pregnancy, uncovering a novel function of maternal liver during pregnancy. Moreover, we revealed an *Ascl1/Igf2* axis in maternal hepatocytes, where Ascl1 normally silences the *Igf2* locus and thereby avoids the exposure of other maternal organs to this growth factor. Others reported that *Ascl1* knockdown resulted in an increase in *Igf2* expression in neuronal cells *in vitro* ^46^. Thus, this axis may be operating in both maternal hepatocytes and neurons. We gained further insight into this axis in that IGF2 was derived from three placental- and fetal-specific promoters of *Igf2* in maternal hepatocytes without *Ascl1*. How *Ascl1* controls the activities of these three promoters of *Igf2* is another question for the future. We believe that this *Ascl1/Igf2* axis, at least in part, underlies *Ascl1*-dependent phenotypes in the maternal compartment. When maternal hepatocyte *Ascl1* is lost, excess IGF2 is produced in, and elaborated from, these cells. Locally, this promotes maternal hepatocyte proliferation and growth. Systemically, it induces overgrowth of maternal pancreas, spleen, and kidney. Through these autocrine and endocrine pathways, IGF2 partially mediates *Ascl1*-dependent phenotypes. It is highly likely that other maternal organs not examined in this report also respond to the null mutation of maternal hepatic *Ascl1*.

Our studies demonstrate that *Ascl1* activation in maternal hepatocytes is required to maintain pregnancy-dependent homeostasis of maternal gut microbiota. It is known that the microbiome shows adaptive changes to accommodate the physiological and immunological alterations of the host during pregnancy ^25, 47, 48^. Here we found that 8 bacteria species in maternal ceca responded to the deficiency of maternal hepatic *Ascl1*. Remarkably, among these bacteria species, *Pseudobutyrivibrio-Roseburia intestinalis* was depleted and *Desulfovibrio oxamicus-vulgaris* was aberrantly appeared. The former is known to be associated with the anti-inflammatory activity of colonic mucosa ^26^, whereas the latter participates in metabolism of various substances such as ammonium, lactate, alcohol, and pyruvate ^27^ Therefore, maternal liver, by activating *Ascl1*, controls these two bacteria populations in maternal ceca, revealing new roles for Ascl1 in maternal liver. Several lines of evidence suggest that the other 6 bacteria species are also important to the health of the host ^26, 27, 49–52^, however, the gestation-dependent functions of these bacteria species remain unclear. Mechanistically, we identified an *Ascl1/Hamp2* axis in maternal hepatocytes as a candidate regulator of maternal gut microbiota. Our data show that *Hamp2* mRNA expression almost fully relies on *Ascl1* activation in maternal hepatocytes. *Hamp2* is produced primarily in the liver, has a strong antimicrobial activity against certain bacteria, and regulates immune responses against bacterial pathogens ^53, 54^. Hence, we assume that this *Ascl1/Hamp2* axis in maternal hepatocytes may partially govern the adaptations of maternal gut microbiota to gestation.

Our studies demonstrate that maternal liver, through activating *Ascl1*, modulates the placenta. This notion is based on our observations that maternal liver *Ascl1* deficiency leads to placenta overgrowth and a change in its function. These phenotypes can be partially interpreted as the consequences of the exposure of the placenta to elevated maternal IGF2. We also observed that hep-*Ascl*^-/-^ placenta transiently reduced its IGF2 production on gestation day 15 and largely diminished its Erk1/2 signaling on gestation day 18 relative to their controls. Thus, the placenta used different mechanisms at distinct stages of pregnancy to defensively respond to increased maternal IGF2, eventually restricting its otherwise further overgrowth. We think that maternal liver activates *Ascl1* to maintain *Igf2* silencing and, by doing this, allows the placenta to appropriately develop and grow without potential maternal inference. It was surprising that the fetal weight was not affected by elevated maternal IGF2. This may suggest that the defense responses of the placenta (reduction in IGF2 production and suppression in Erk1/2 activity) to elevated maternal IGF2 is so strong that the potential stimulatory effect of this potent hormone on fetal growth is fully blocked. However, we did observe severely impaired postnatal growth of the offspring born from hep-*Ascl1*^-/-^ dam. It is a well-established concept that maternal problems generate long-term adverse effects on the health of the next generation ^55–57^. Hence, we believe that *Ascl1* deficiency-caused dysregulation of maternal adaptations and placental abnormality together impair the postnatal growth of the offspring. This model warrants further investigations.

In summary, we demonstrate that, as pregnancy advances, maternal hepatocytes induce expression of *Ascl1* to alter their transcriptomes. Via this mechanism, maternal liver systematically coordinates adaptations in the maternal compartment and allows for optimal placental development and growth, collectively ensuring the health of both the mother and her infant during pregnancy and postnatally.

## Author contributions

Conceptualization: JL and GD. Investigation: JL, VG, SN, and HJ. Formal analysis: JL, VG, and HJ. Data curation: JL, VG, and HJ. Writing: JL and GD. Resources: GD.

## Abbreviations

Ascl1: achaete-scute homolog-like 1
IGF2: insulin-like growth factor 2
AST: aspartate aminotransferase
ALT: alanine aminotransferase
PL: placental lactogen
GD: gestation day.

---

**Supplemental Fig. 1.**
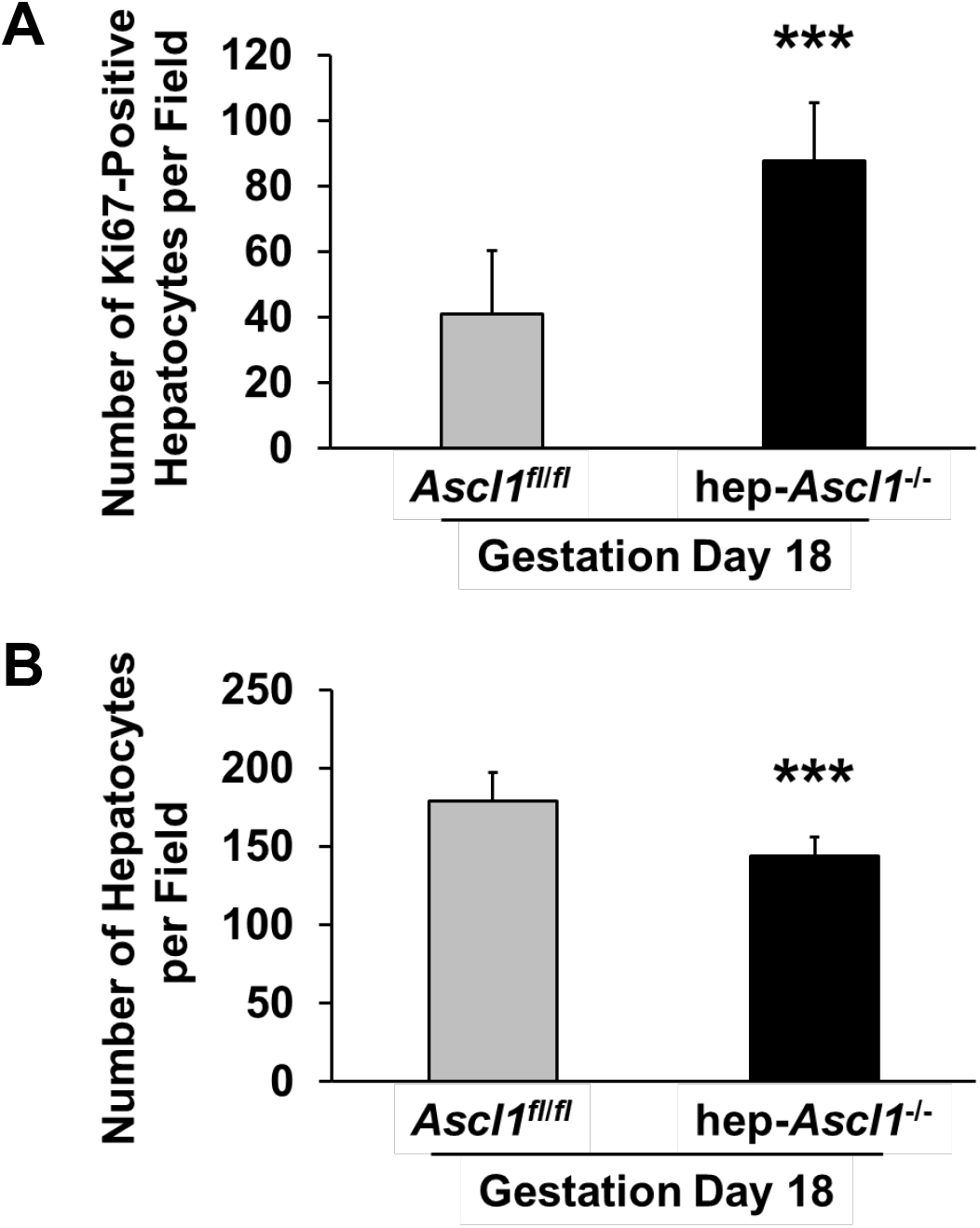
Maternal hepatocyte-specific *Ascl1* knockout results in increased maternal hepatocyte proliferation and hypertrophy. Livers were collected from nonpregnant (NP) and gestation day (GD) 18 *Ascl1^fl/fl^* and *hep-Ascl1^-/-^* mice. Liver sections were subjected to **(A)** Ki67 and **(B)** β-Catenin staining. Ki67- and β-Catenin-positive hepatocytes were counted in five random fields of view (X200 magnification) and presented as the means ± SD (n = 3-6 and n=7 for each group, respectively). ***, *P* < 0.001, between *Ascl1*^fl/fl^ and hep-*Ascl1*^-/-^ mice.

**Supplemental Fig. 2.**
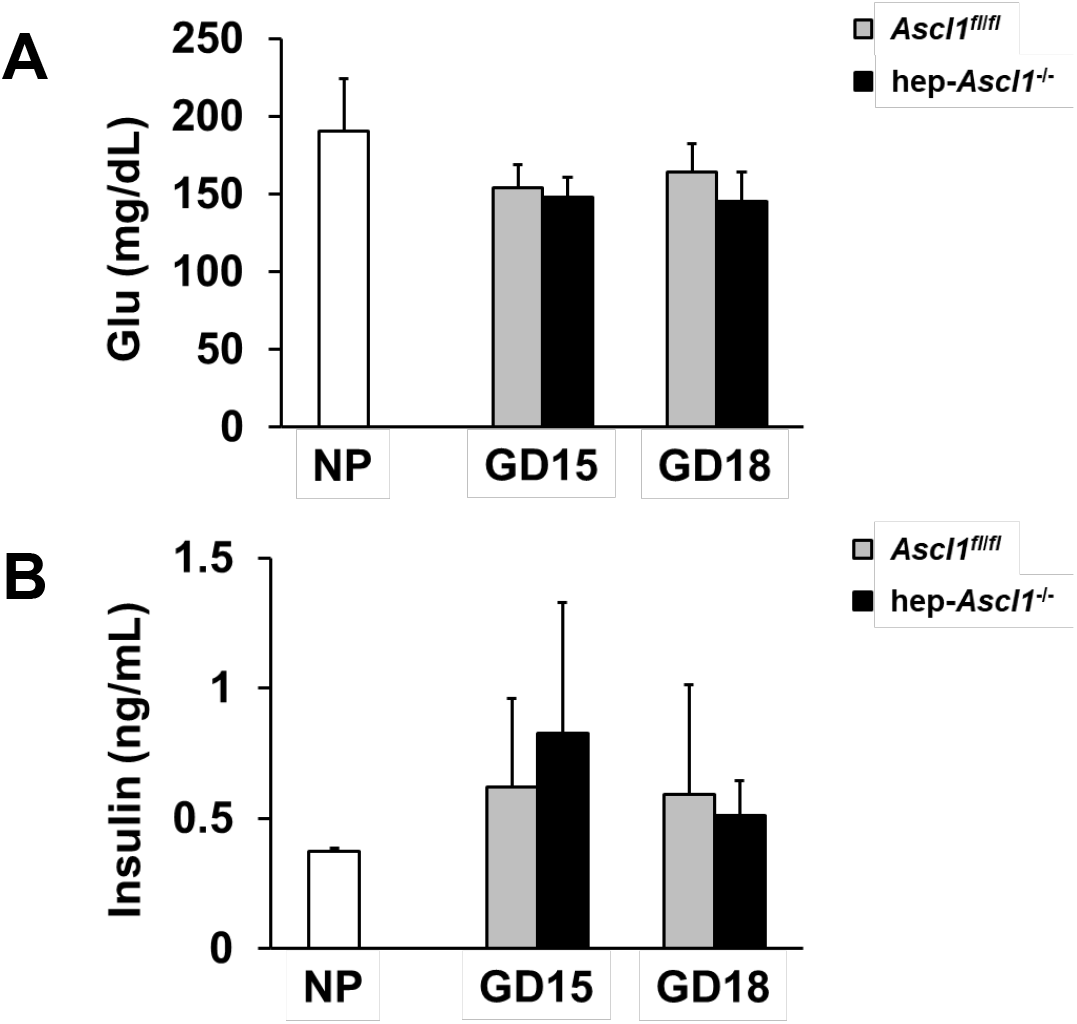
Maternal hepatocyte-specific *Ascl1* knockout does not affect the concentrations of glucose and insulin in maternal blood. Serums were collected from nonpregnant (NP) and gestation days (GD) 15 and 18 *Ascl1^fl/fl^* and *hep-Ascl1^-/-^* mice. Serum **(A)** glucose and **(B)** insulin are presented as means ± SD (n = 3-6). *Ascl1*, achaete-scute homolog 1; Glu, glucose.

**Supplemental Fig. 3.**
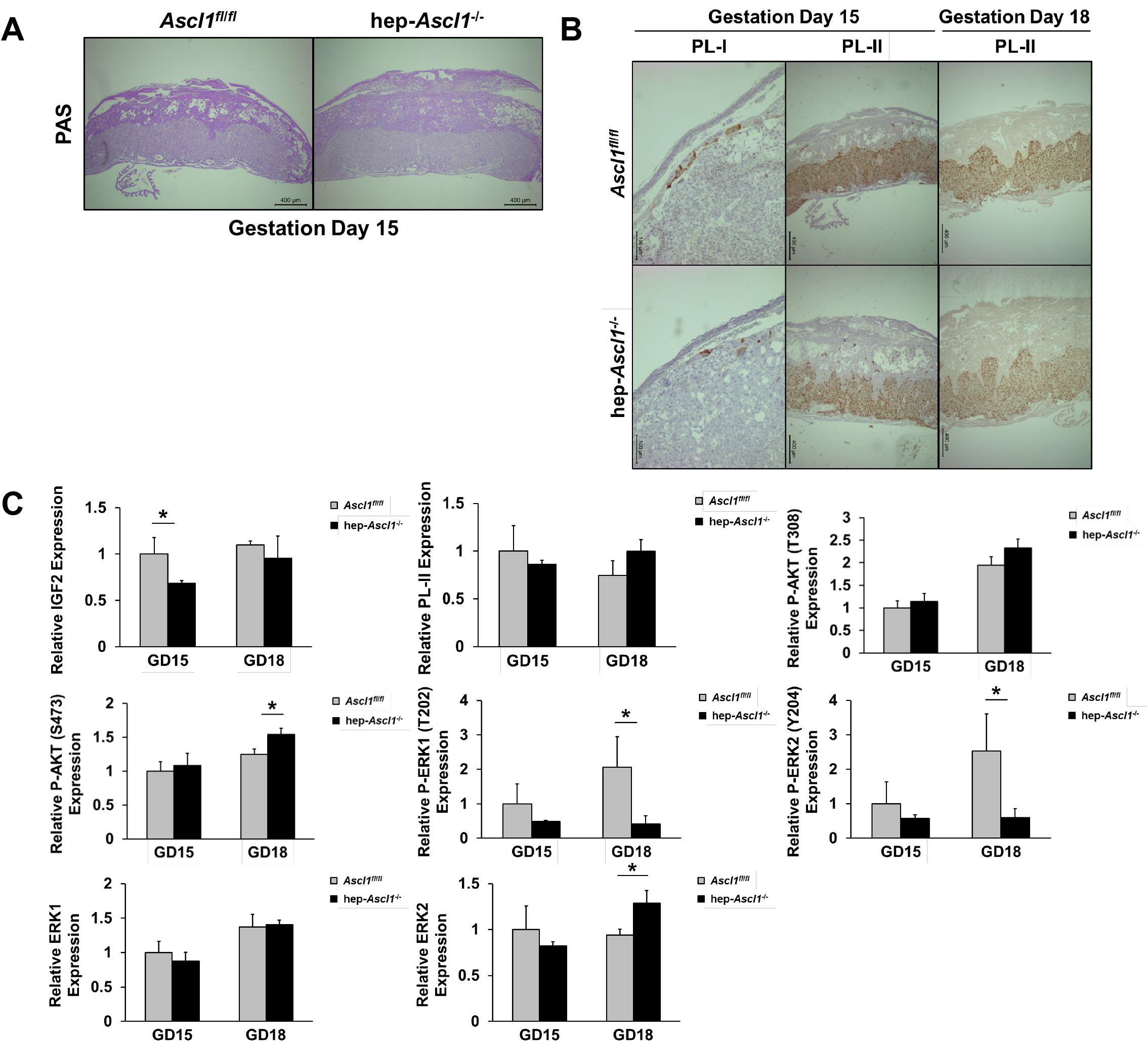
Maternal hepatocyte-specific *Ascl1* knockout partially influences placental function. Placentas were collected from gestation days (GD) 15 and 18 *Ascl1^fl/fl^* and *hep-Ascl1^-/-^* mice. **(A)** Frozen placental sections underwent periodic acid-Schiff (PAS) staining. Glycogen is stained red to purple. **(B)** Frozen placental sections were subjected to *PL-I* and *PL-II in situ* hybridization staining using RNAscope 2.5 HD Assay-BROWN kit. The *PL-I* and *PL-II* mRNAs are stained dark brown. **(C)** Quantification of relative intensity of Western blotting signals from **Figure 6D**. Data are presented as the mean fold changes relative to GD15 *Ascl1^fl/fl^* mice (± SD; n = 3 for each group). *, *P* < 0.05, between *Ascl1^fl/fl^* and *hep-Ascl1^-/-^* mice. AKT, total protein kinase B; *Ascl1*, achaete-scute homolog 1; ERK1/2, extracellular signal-regulated kinase 1/2; GAPDH, glyceraldehyde 3-phosphate dehydrogenase; IGF2, insulin-like growth factor; PL-II, placental lactogen II.

**Supplemental Fig. 4.**
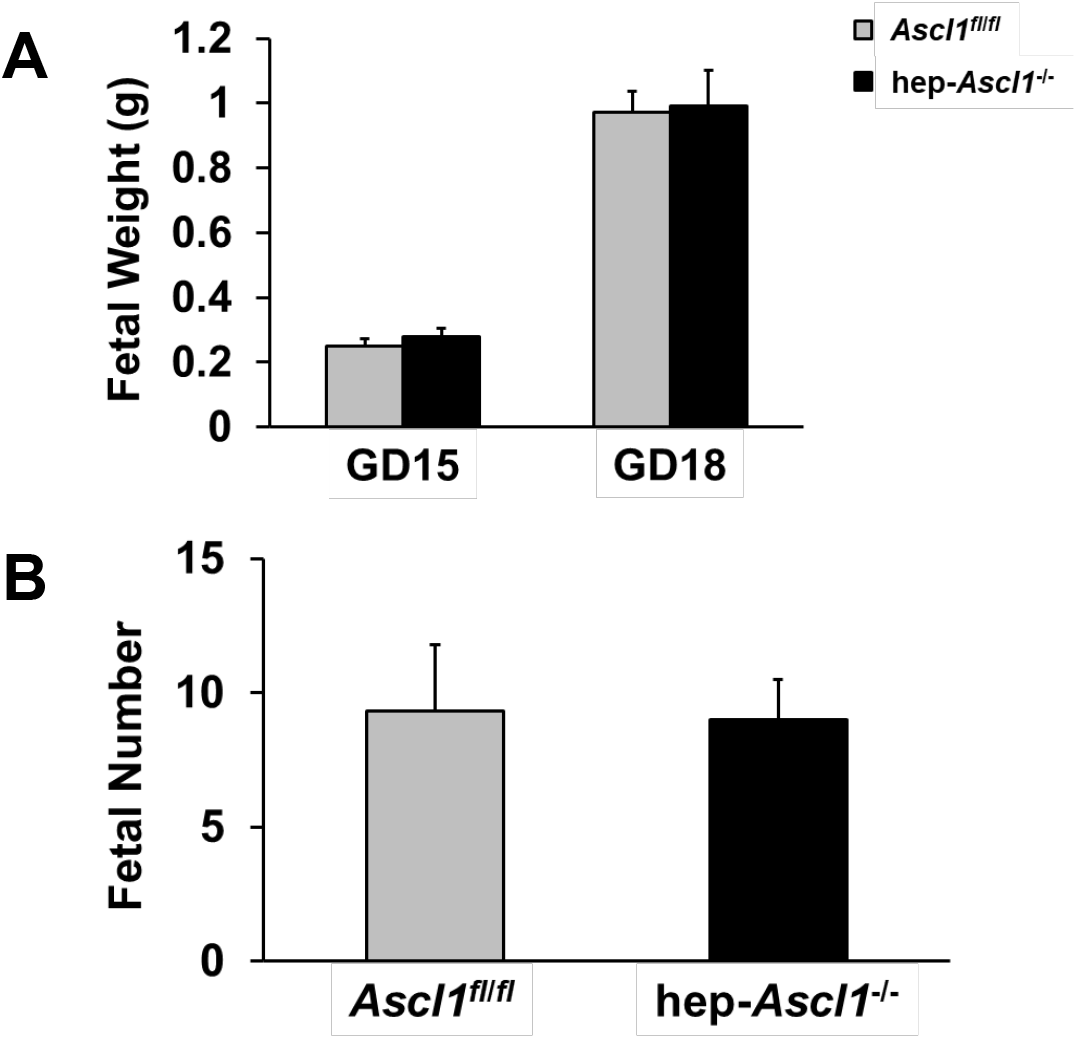
Maternal hepatocyte-specific *Ascl1* knockout does not affect fetal weight and numbers. Fetuses were weighed from gestation days (GD) 15 and 18 *Ascl1^fl/fl^* and hep-*Ascl1*^-/-^ mice. **(A)** Fetal weight and **(B)** numbers are presented as means ± SD (n = 13-31). *Ascl1*, achaete-scute homolog 1.

**Supplemental Table S1.**
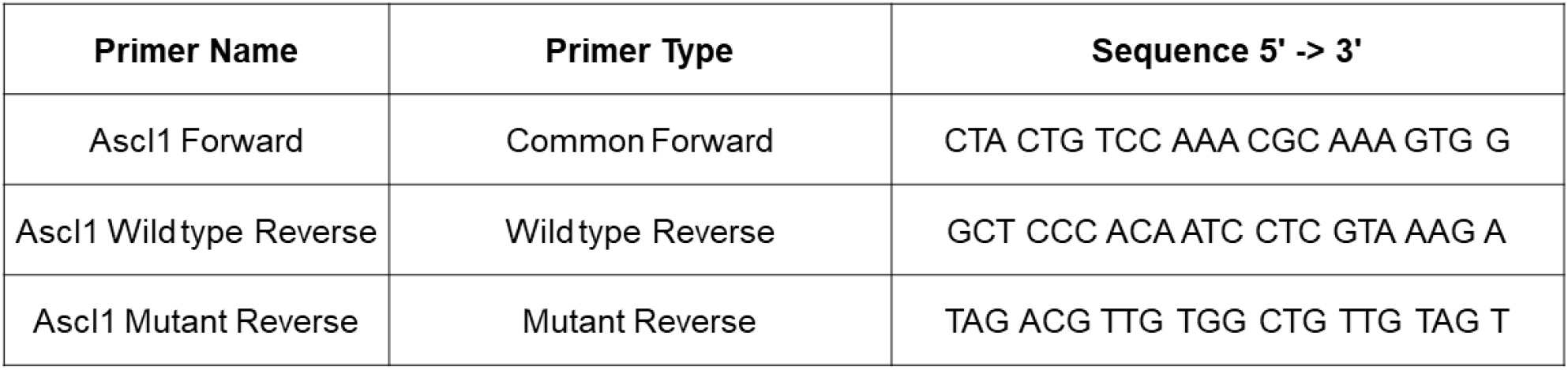

**Supplemental Table S2.**

| Gene | Assay ID | Catalog No. |
| --- | --- | --- |
| Ascl1 | Mm03058063_m1 | 4453320 |
| Igf2 | Mm00439564_m1 | 4331182 |
| Hamp2 | Mm00842044_g1 | 4331182 |
| 18S | Mm03928990_g1 | 4453320 |

**Supplemental Table S3.**
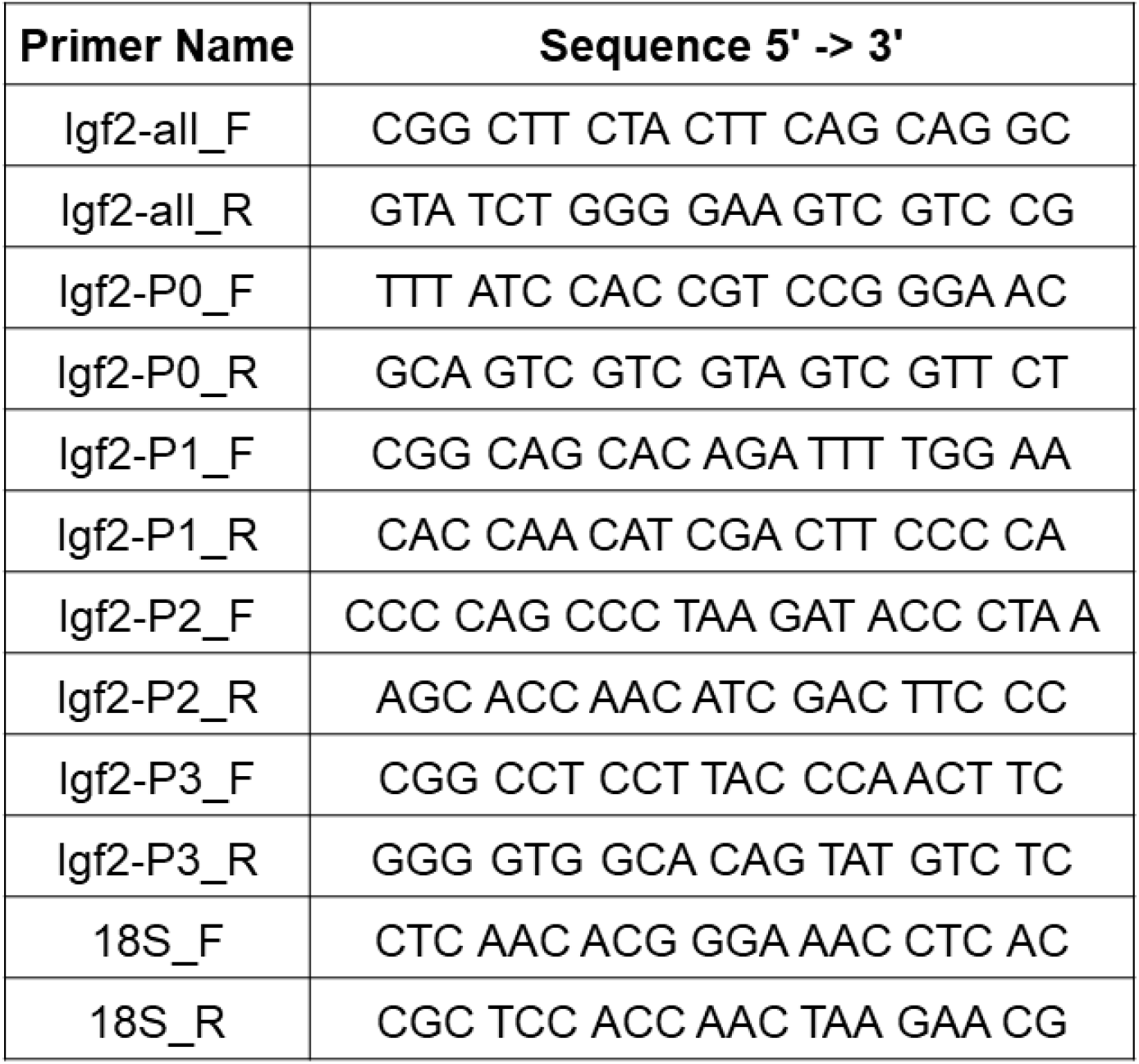

